# Using Topic Modeling via Non-negative Matrix Factorization to Identify Relationships between Genetic Variants and Disease Phenotypes: A Case Study of Lipoprotein(a) (*LPA*)

**DOI:** 10.1101/335745

**Authors:** Juan Zhao, QiPing Feng, Patrick Wu, Jeremy L. Warner, Joshua C. Denny, Wei-Qi Wei

## Abstract

Genome-wide and phenome-wide association studies are commonly used to identify important relationships between genetic variants and phenotypes. Most of these studies have treated diseases as independent variables and suffered from heavy multiple adjustment burdens due to the large number of genetic variants and disease phenotypes. In this study, we propose using topic modeling via non-negative matrix factorization (NMF) for identifying associations between disease phenotypes and genetic variants. Topic modeling is an unsupervised machine learning approach that can be used to learn the semantic patterns from electronic health record data. We chose rs10455872 in *LPA* as the predictor since it has been shown to be associated with increased risk of hyperlipidemia and cardiovascular diseases (CVD). Using data of 12,759 individuals from the biobank at Vanderbilt University Medical Center, we trained a topic model using NMF from 1,853 distinct phecodes extracted from the cohort’s electronic health records and generated six topics. We quantified their associations with rs10455872 in *LPA*. Topics indicating CVD had positive correlations with rs10455872 *(P* < 0.001), replicating a previous finding. We also identified a negative correlation between *LPA* and a topic representing lung cancer (*P* < 0.001). Our results demonstrate the applicability of topic modeling in exploring the relationship between the genome and clinical diseases.

**Author summary:** Identifying the clinical associations of genetic variants remains crucial in understanding how the human genome modulates disease risk. Traditional phenome-wide association studies consider each disease phenotype as an independent variable, however, diseases often present as complex clusters of comorbid conditions. In this study, we propose using topic modeling to model electronic health record data as a mixture of topics (e.g., disease clusters or relevant comorbidities) and testing associations between topics and genetic variants. Our results demonstrated the feasibility of using topic modeling to replicate and discover novel associations between the human genome and clinical diseases.

## Introduction

Elucidating associations between genetic variants and human diseases creates new avenues for disease prevention and enables more precise treatment of diseases [1,2]. During the past two decades, genetic studies have uncovered thousands of genetic variants that influence risk for disease phenotypes [3], e.g., the discovery of a variant in proprotein convertase subtilisin/kexin type 9 (*PCSK9[4]*) associated with low plasma low-density lipoprotein, which led to a new therapeutic drug class that was approved by the US Food and Drug Administration in 2015. Many of these discoveries come from large-scale association analyses. The two most notable approaches are genome-wide (GWAS) and phenome-wide association studies (PheWAS) [2, 5]. For a given phenotype, GWAS scans hundreds of thousands to millions of single nucleotide polymorphisms (SNPs) across the genome in a hypothesis-free approach. PheWAS, on the contrary, analyzes thousands of disease phenotypes compared to a single SNP. In a GWAS, the outcome variable is a disease phenotype and predictor variables are SNPs. While in a PheWAS, the outcome variable is a SNP and predictor variables are disease phenotypes.

Association analyses test a large number of predictor variables at one time and assume that each variable has an independent effect. However, diseases often occur together as a group of comorbidities, e.g. hyperlipidemia (HLD) and cardiovascular diseases (CVDs). Conventional association analyses may not capture the inter-connections among variables such as phenotypes and thus may not be sensitive enough to identify important genotype-phenotype relationships. Moreover, association analyses also face the challenge of scaling to an increasing number of phenotypes. Previously, we have described a “networked PheWAS” approach which can address interconnectivity but still requires a degree of supervised interpretation [6].

This study aimed to test the feasibility of topic modeling for identifying relationships between genetic variants and disease phenotypes. Topic modeling is an unsupervised machine learning method that was initially introduced as a text mining technique [7]. It has been demonstrated to extract latent topics or themes from documents, aiding in the understanding of large amounts of data [8]. Compared to traditional clustering approaches such as K-means clustering that partitions a collection of documents into several disjoint clusters (i.e., topics) based on a similarity measure, topic modeling assigns a document to multiple clusters with different scores. Therefore, each document is characterized by one or more topics. In addition to its wide adoption in the text mining field, topic modeling has achieved many successes in computer vision and biomedical science. Recently, a few groups have used this approach to analyze electronic health records (EHRs) [9,10] and genetic data to capture the characteristic of data [11,12].

We hypothesized that topic modeling would be useful in replicating known findings and uncovering previously unidentified relationships between genetic variants and disease phenotypes. To test this hypothesis, we used topic modeling via non-negative matrix factorization (NMF) [13,14] to identify latent topics (e.g. disease clusters or relevant comorbidities) from EHR data. We then tested associations between the EHR-derived topics and a *LPA* SNP (rs10455872). We chose the SNP because previous studies have shown that high-levels of the *LPA* product (Lp(a)) is associated with increased risks of developing HLD and CVD [15]. Specifically, rs10455872, as a single variant, explains 20-30% of the variation in circulating Lp(a) levels, which makes it an ideal candidate for this study [16].

## Results

We applied a topic modeling algorithm using NMF on the dataset of 12,759 individuals and obtained six potentially meaningful topics from the EHRs (Fig 1). The learned topics (i.e., clusters of disease phenotypes) were consistent with the comorbidities associated with the phenotypes most prevalent in the cohort. For example, topic #0 represented diseases of respiratory failure, topic #2 defined diseases related to CVD (e.g., HLD, hypertension, and chronic ischemic heart disease), topic #3 represented phenotypes relevant to lung cancer and its treatment, topic #4 was related to diabetes and its comorbidities; and topic #5 was related to liver disease and its sequelae.

**Fig 1.**
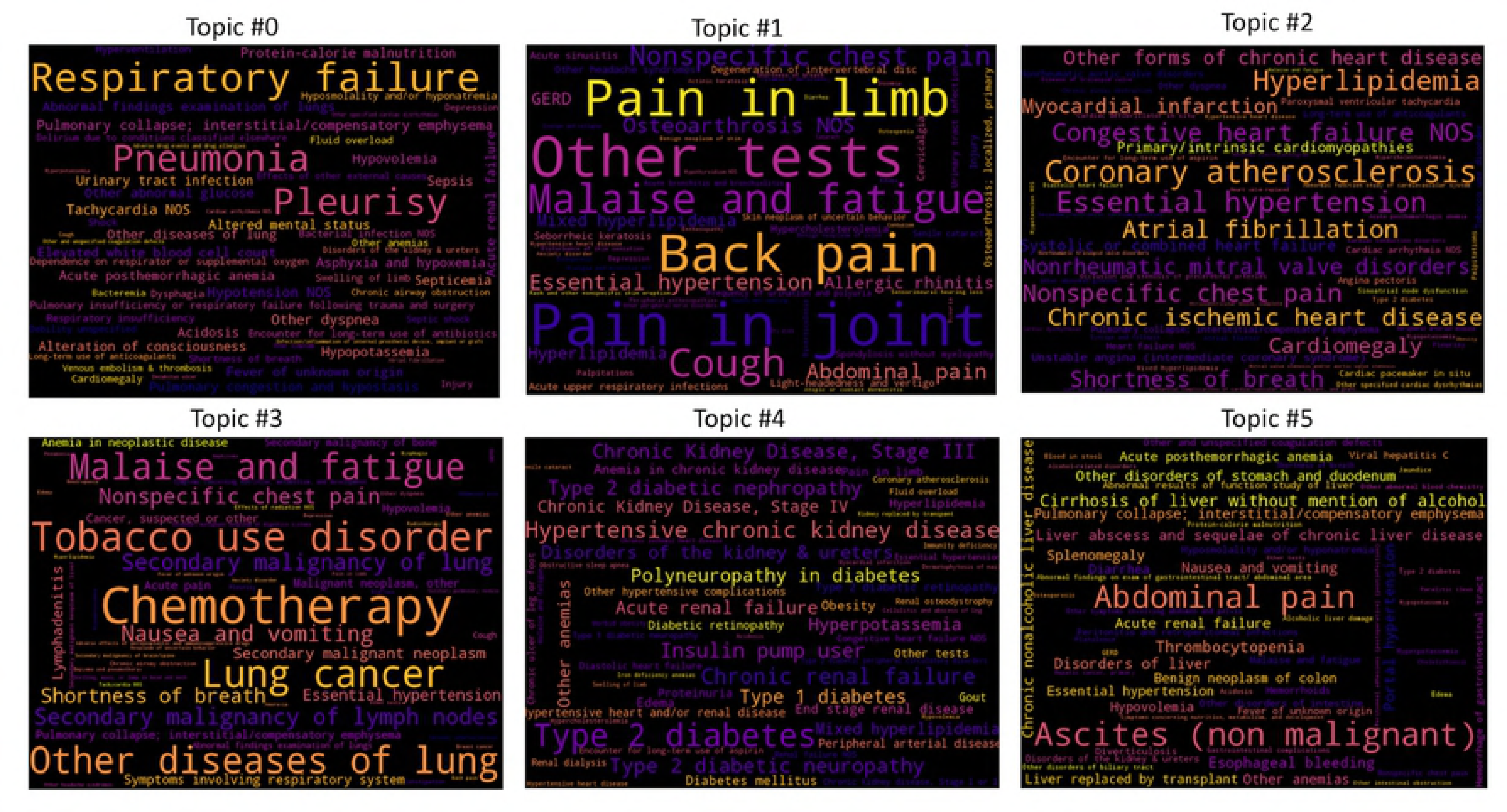
Word clouds for six topics. The size of the words (phecode) in each cloud indicate the weights of the phenotypes on the topic. Phenotypes with larger-sized words had greater influence on the topic compared to phenotypes with smaller-sized words. For each word cloud, we listed the top 60 words to provide a better visual presentation of what each topic represents.

Fig 2 shows the distribution of the numbers of topics in the cohort. Topic #2 was the most prevalent (33%) topic in the cohort. Topics #1 and #3 were the second and third most prevalent topics in the cohort.

**Fig 2.**
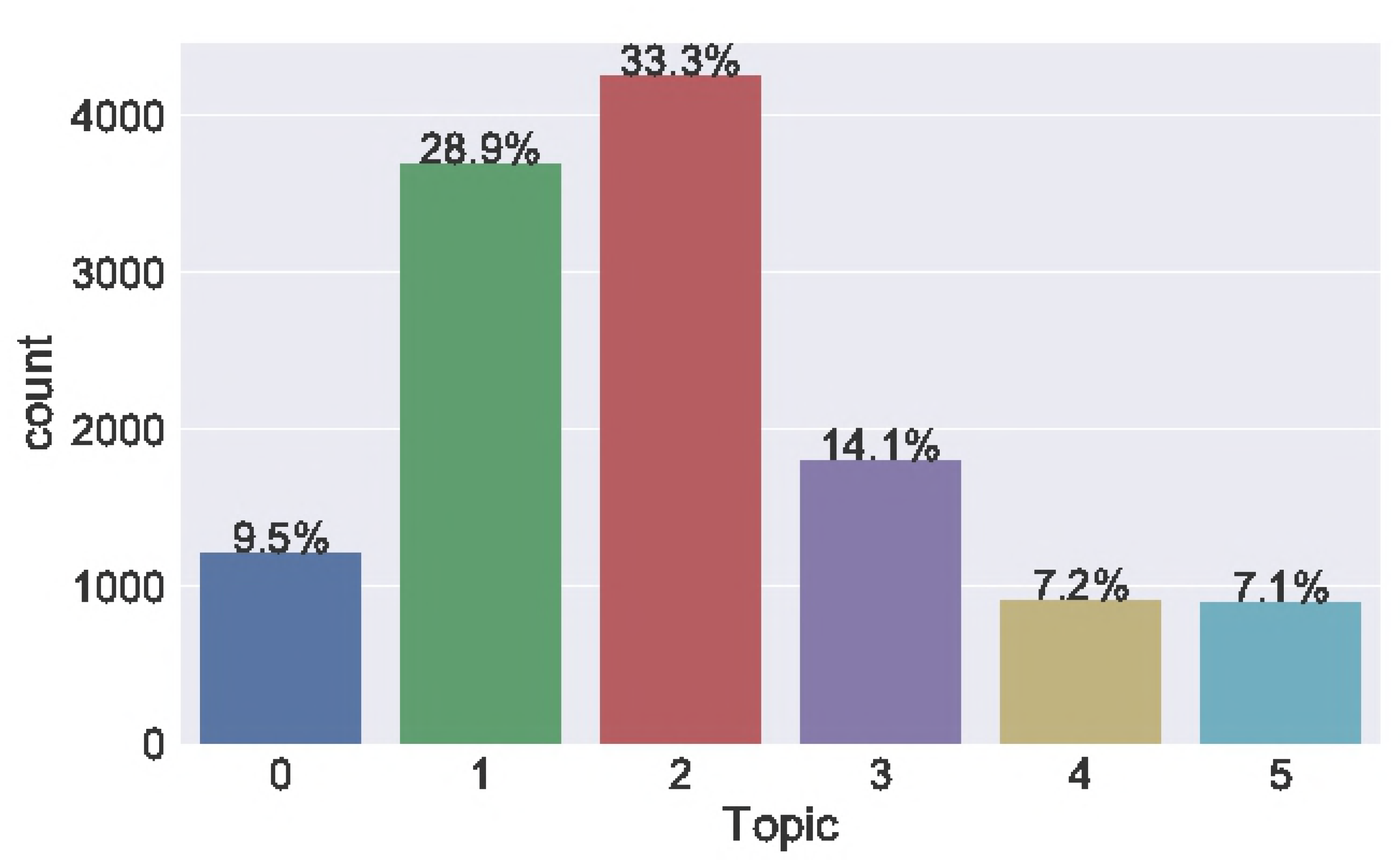
Topic distribution in the cohort. To visualize the prevalence of each topic in the cohort, we assigned an individual to the topic with the maximum score.

We also used t-Distributed Stochastic Neighbor Embedding (t-SNE) [17] to transform the individual-phenotypes matrix (*W*) into a 2-dimensional (2D) space to visualize the quality of topic modeling (Fig 3). Each data point in the figure corresponds to one individual. We labeled each individual with the assigned topic.

**Fig 3.**
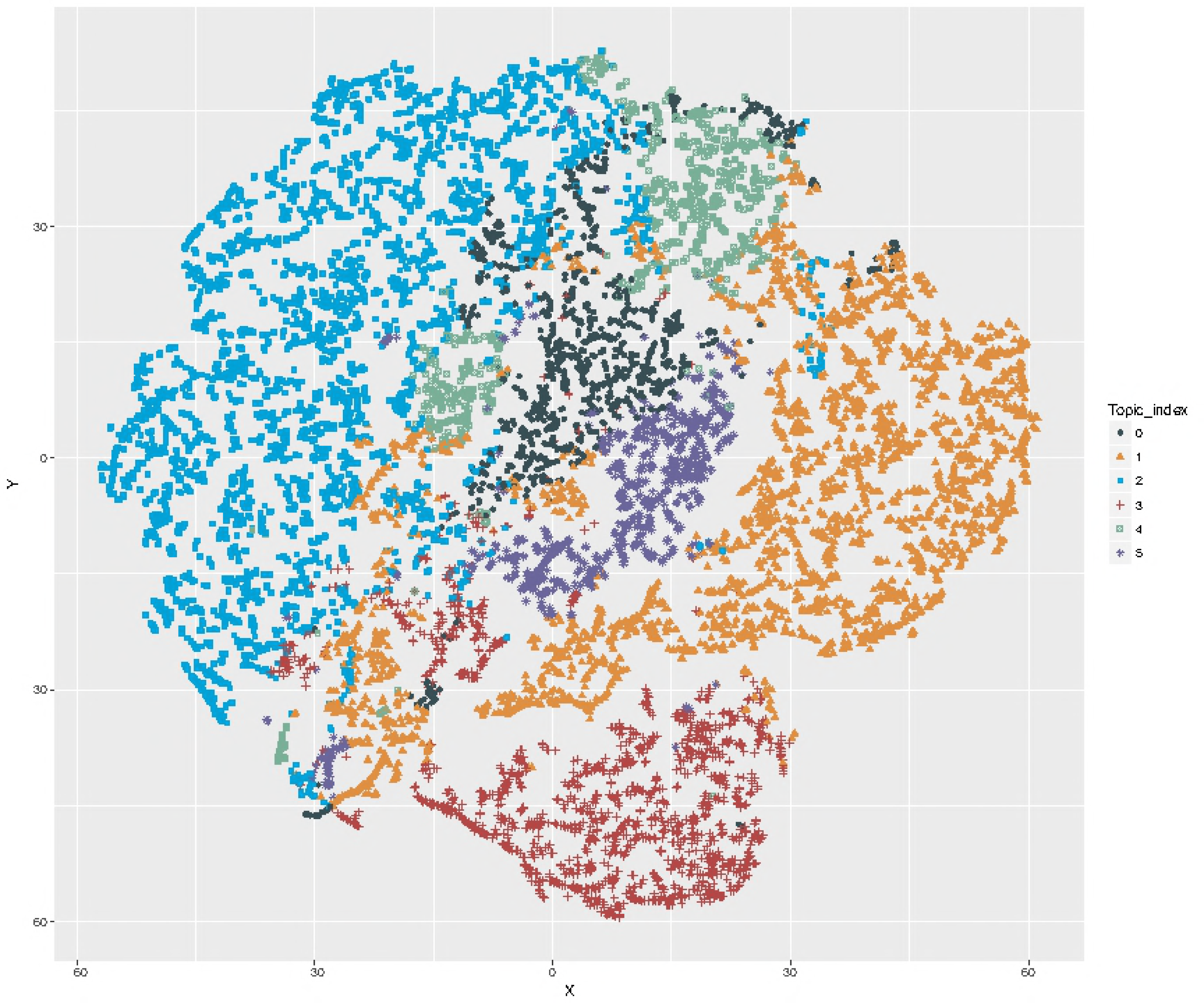
t-SNE plot of visualizing the patient clusters in a projected 2D metric map.

The perplexity was set to 30. We used PCA initialization as it is more globally stable. Each point represents an individual. Topic #2 contains the most individuals in the cohort.

We then applied the Pearson correlation coefficient (PCC) to examine the association between each topic and rs10455872. Statistical test results suggest that topic #2 and #3 were significantly associated with rs10455872 (Table 1). Topic #2, a group of lipid and cardiovascular diseases, had a positive correlation with rs10455872 (r=0.072, p=5.8e-16). We also found that topic #3, a group of phenotypes relevant to lung cancer, had a negative correlation with rs10455872 (r=-0.039, p=8.5e-6). Although the *r* coefficient is weaker than the topic#2, these correlations are highly statistically significant.

**Table 1.**
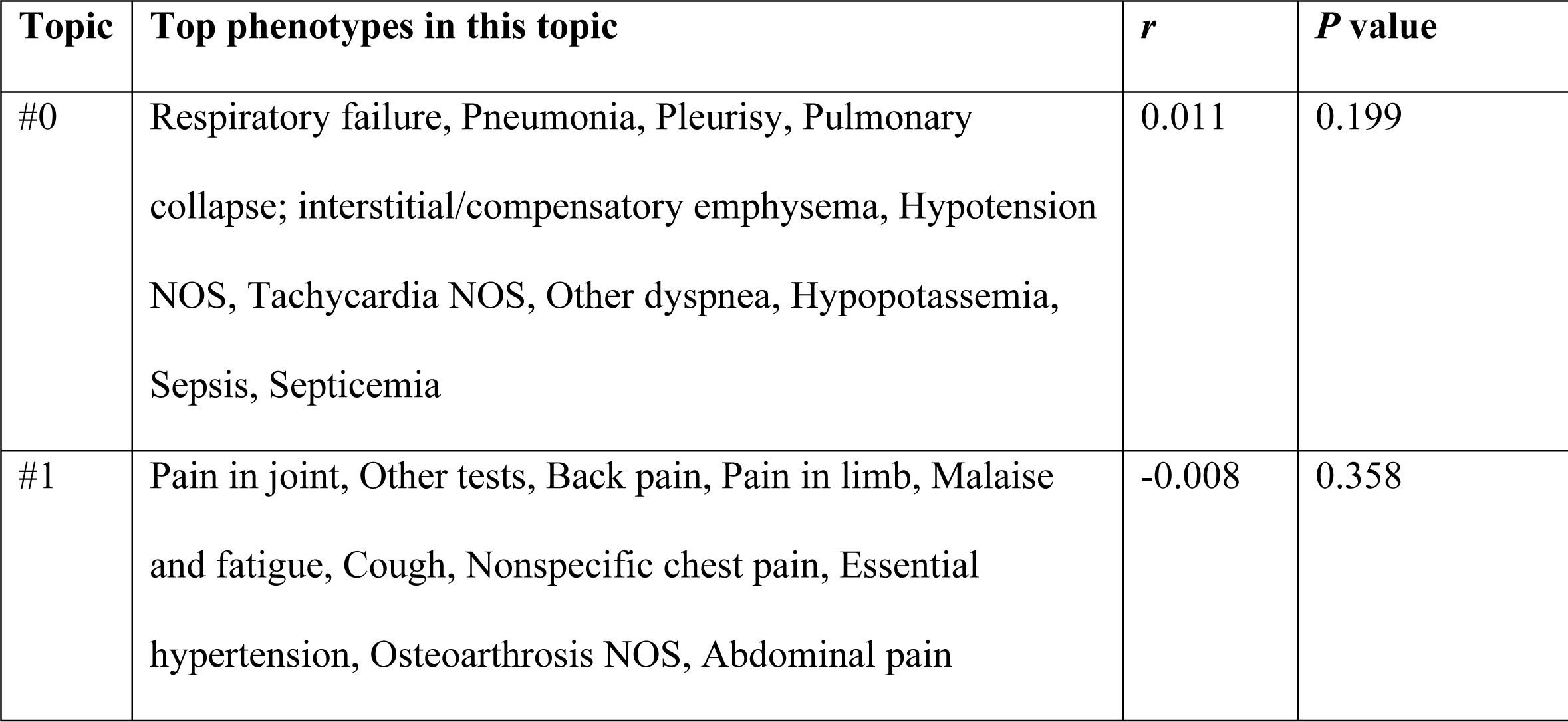
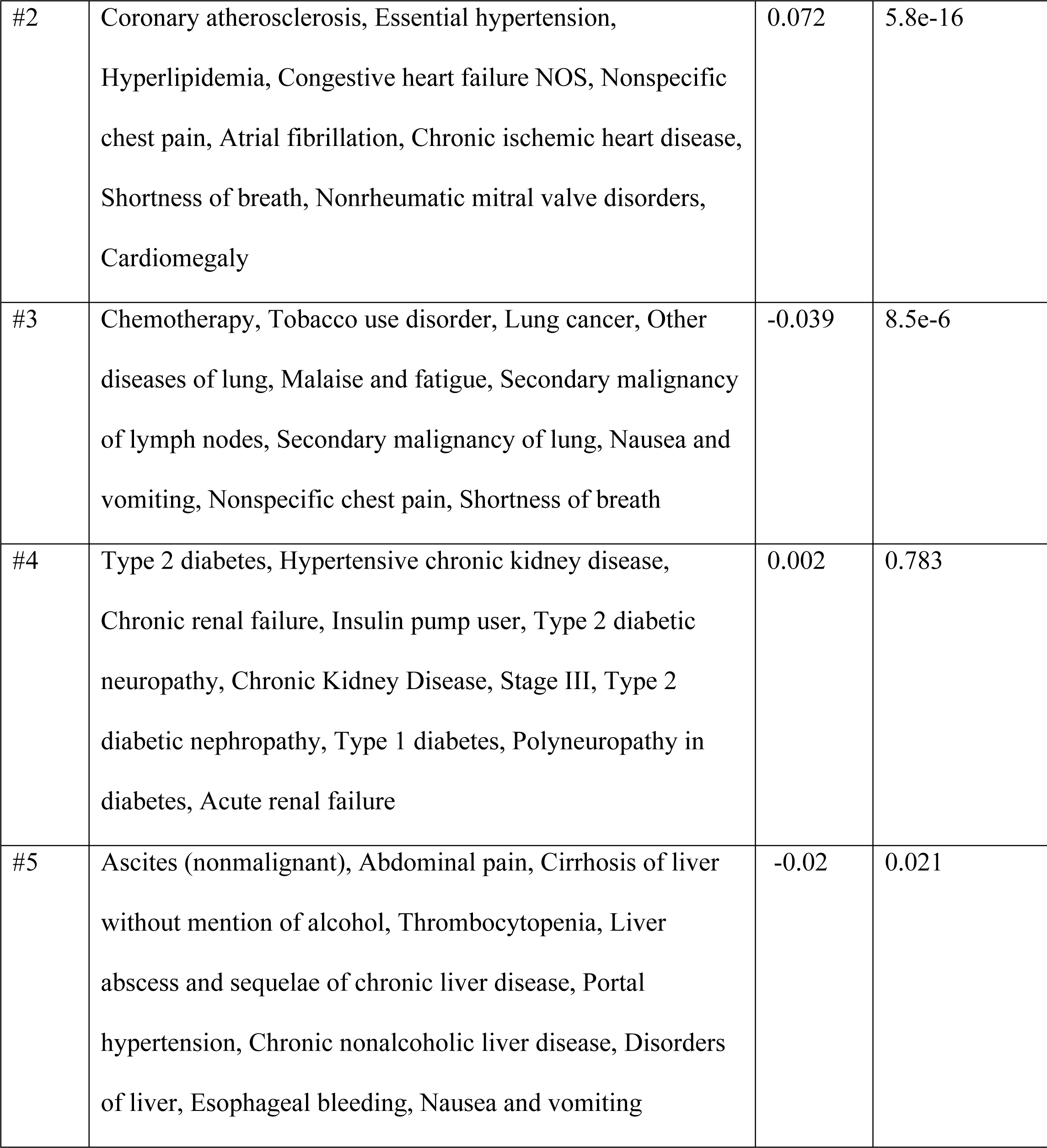
Pearson correlation between LPA variant for each topic

## Discussion

Topic modeling has been widely used in the field of text mining. In this paper, we applied this technique to explore associations between disease phenotypes and genetic variants. We assumed that some disease phenotypes found simultaneously in a large EHR have correlated semantic meanings and thus can be learned as topics. We examined the associations between a *LPA* variant (rs10455872) and the six topics derived from EHRs. We observed the expected association between rs10455872 and a topic representing CVD/HLD. We also found a novel association, as of this writing [18], between the *LPA* variant and a lung cancer topic.

The *LPA* gene encodes lipoprotein (a), a major component of the Lp(a) particle.

Individuals with elevated Lp(a) levels are more likely to develop CVD compared to those with normal or low Lp(a) level [16,19]. Approximately 70% of Lp(a) variation can be attributed to variants at the *LPA* locus [20–22], and rs10455872 alone explains ∼25% variation in circulating Lp(a) levels [16]. Further, a previous genetic study suggested that *LPA* variants were strong predictors for CVD risk [16]. In a more recent study of >10,000 patients taking statins, our group found that rs10455872 predicted residual CVD risk while on lipid-lowering treatment [23]. This study’s finding of a significant association between rs10455872 and the CVD/HLD topic demonstrates the feasibility of topic modeling as a critical tool for uncovering genotype-phenotype relationships.

We also observed a negative correlation between the *LPA* variant and the cancer/lung cancer topic, i.e., possessing this variant is protective. Previous epidemiological studies have reported that individuals with low Lp(a) levels had increased risk of all-cause and cancer-related mortality [24]. Mieno et al. found that hypolipoproteinemia(a) is a risk factor for cancer except for lung cancer. Nevertheless, there are few reports on a relationship between cancer and *LPA* polymorphism or expression levels. Our previous PheWAS analysis of a separate cohort identified an association between rs10455872 and cancer diagnosis code with borderline significance [23]. To further explore this association between rs10455872 and the cancer/lung cancer topic, we queried gene2pheno (https://imlab.shinyapps.io/gene2pheno_ukb_neale/), which is a publicly available database for testing associations between predicted gene expression levels and phenotypes using data from the UK Biobank. Genetically predicted LPA expression levels were associated with death from T cell lymphomas (p=6.9 e-5, Underlying (primary) cause of death: ICD10: C84.5 Other and unspecified T-cell lymphomas). Given that lung cancer is strongly mediated by environmental exposure and that tobacco use disorder was also part of topic #4, it is possible that the SNP is a marker for propensity to smoking, e.g., similar to what was shown for rs16969968 [25]. Further genetic and epidemiological studies are needed to elucidate the relationship between Lp(a) levels and cancer incidence.

Topic modeling approaches require pre-specification of the number of topics. In this study, we set *k*=6, because we aimed to capture the most prevalent diseases such as CVD and to quantify the association. Increasing *k* allows the quantification of associations between genetic variants and rare diseases but risks fracturing common phenotype clusters. It can be seen that (Fig 3), except for topic #4 (diabetes), the learned topics formed distinct clusters, indicating a good quality of topic modeling. Some of points in topic #4 (diabetes) were close with topic #2 (CVD), which was expected, because type II diabetes is an important risk factor that increases the risk of developing CVD. Compared to the other topics, #1 (Pain), #2 (CVD), and #3 (Lung Cancer) have more concentrated clusters.

For optimal selection of *k*, common approaches have used different values of *k* to look at the error in optimization and selected the best value by having domain experts review the topics to identify which set of topics are most meaningful, and have estimated *k* using singular value decomposition (SVD) to look at the decay of singular values [26–28]. To provide evidence for the stabilities of our results, we also set different numbers of topics *k* = 10, 20, 30 (Supplementary Table 1) and examined the PCC. Results were consistent with topics at *k*=6.

In summary, unlike traditional PheWAS that have treated each disease phenotype as a distinct variable, topic modeling via NMF generates more abstract latent factors from disease phenotypes and significantly reduces the number of multiple tests. Our results demonstrate the power of topic modeling in the detection of comorbidities and previously unexplored genotype-phenotype relationships among a large cohort.

## Limitations

There are several limitations in this study. First, we tested only one genetic variant in one gene. Rs10455872 explains approximate 25% change in circulating Lp(a) levels according to previous studies; however, it would be interesting to generate a genetic risk score for Lp(a) levels and test its association with disease phenotypes in the future. Second, we used a binary value to indicate if an individual had a diagnosis code. A method accounting for disease severity (e.g., counts of diagnosis codes) could be used in future studies. Finally, the current study was limited to using billing codes to phenotype individuals. We did not include other information, e.g. laboratory test and medications, to assign more accurate phenotypes. This problem can be solved in the future using more sophisticated “deep” phenotyping methods that include more features from EHRs.

## Materials and methods

### Study cohort

We used data from BioVU, the de-identified DNA biobank at Vanderbilt University Medical Center (VUMC), to conduct this study. BioVU contains DNA samples from >250,000 individuals that are linked with their de-identified EHRs, including diagnostic and procedure codes, clinical notes, laboratory values and medications. We identified 12,759 adult individuals of European ancestry (F/M: 6,018/6,741; age: 70.3±12.3) who had both EHRs and genotyped data of rs10455872 available.

### rs10455872 Genotyping

We extracted each individual’s rs10455872 information from existing genotyped data. All genotyping was previously conducted using commercially available genome-wide SNP arrays with quality control criteria for variants followed by a standard imputation process using 1000 Genomes Project allele frequency estimates.

Among the cohort of 12,759 individuals, we observed 85.2% AA, 14.2% AG, 6.1% GG. The minor allele frequency (MAF) of the rs10455872 G allele is 7.7% in our cohort, consistent with the 7% MAF in the European population [29]. We used 0, 1, 2 to represent the number of *LPA* rs10455872 G alleles that an individual carry.

### Disease Phenotypes

Following established protocols used in past studies [30], we grouped each individual’s ICD-9-CM (International Classification of Disease, 9^th^ edition) codes into disease phecodes. There were 1853 phecodes present in the 12,759 individuals. For each phecode, we labeled individuals without the phecode with a ‘0’ and those with the phecode with a ‘1’.

We applied a topic modeling algorithm using NMF on the dataset of 12,759 individuals to learn potentially meaningful topics from the EHRs. Then, we quantified the association between each learned topic with rs10455872 using PCC. The workflow of this experiment is demonstrated in Fig 4.

**Fig 4.**
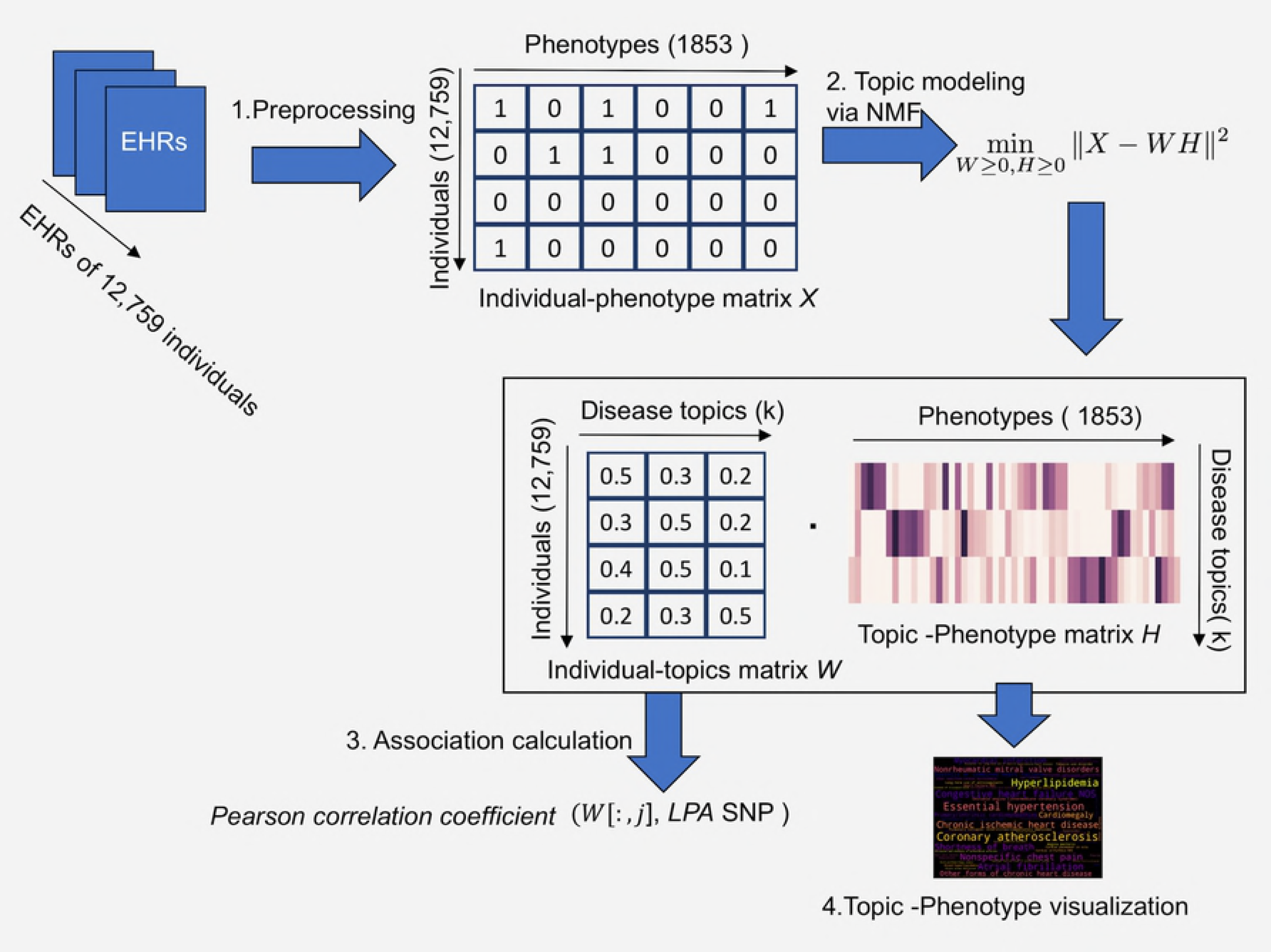
Illustration of topic modeling on EHRs using NMF.

### Topic modeling via Non-negative Matrix Factorization (NMF)

We used NMF as our topic modeling approach. NMF is a low-rank matrix approximation algorithm that has been widely used for feature reduction for high-dimensional data. The assumption is that given a large and sparse matrix *X* of size *n× m* representing a collection of *n* high dimensional data points in *R*^*m*^. *X* is low rank which means that most data points can be approximately represented by a linear combination of a small set of *k* basis vectors *H* _∈_ *R*^*k × n*^. The linear combination is a coefficients matrix *W* _∈_ *R*^*n × k*^ providing a lower-dimensional encoding for *X*, which result in a feature reduction for a high-dimensional data. Since NMF restricts the *X* non-negative and enforces the *H* and *W* to be also non-negative, NMF has good interpretability and has been commonly used as a topic modeling approach in text mining.

We considered each individual’s EHR as one document, and each document was described by disease phenotypes represented by the phecodes (Fig 1). Since we had 12,759 (*n)* individuals’ EHRs and 1,853 (*m)* unique phecodes, we used matrix *X* _∈_ *R*^*n × m*^ to represent the input data, where each row of *X* represented an individual, and each column of *X* was a phecode. The entry of the matrix *X*_*ij* ∈_ *X* was a binary value (0 or 1) indicating whether *i*th individual had the *j*th phecode. This representation is similar with the bag-of-word model, where each document is associated with a set of words, and word ordering in the documents does not matter.

Given that non-negative input matrix _∈_ *R*^*n × m*^, and an expected number of topics *k*_≤_ *m*, NMF generates two matrices, *W* _∈_ *R*^*n × k*^, and *H* _∈_ *R*^*k× m*^. Both *W* and *H* are non-negative entries, such that

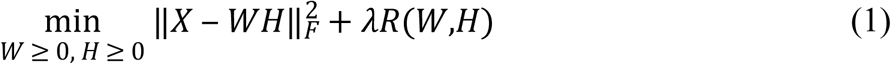

*H*(*k*,:) is a latent topic – phenotype matrix. Specifically, each row of *H* corresponds to a disease topic, and each topic is represented by a set of relevant phenotypes that co-occurred in several individuals’ EHRs, with specific cores indicating their relevance to this topic. Through *H*(*k*,:), we extracted a semantic meaning of each topic, e.g. what kind of diseases or comorbidities are represented by the topic.

*W*(*i,k*) is an individual-topic matrix. Each row of *W* corresponds to an individual’s score on each topic that indicates the diseases and comorbidities carried by the individual. An individual that has a large score for a disease topic indicates that there a higher probability for an association between individual and the topic. *W*(*i,k*) is then used for the association calculation between the topics and rs10455872, which is described further below.

*R*(*W, H*) is the regularization term that combines L1 and L2 norms, which is defined as:

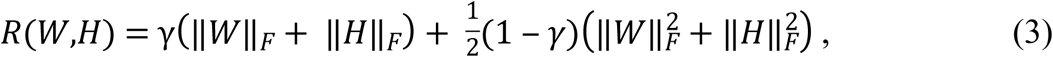

where *γ* is the ratio for L1 penalty. Adding the regularization term is necessary for balancing the sparsity of the topics, meaning that an individual may have several topics at the same time. Moreover, addition of the regularization term minimizes the effect of outliers on the model.

### Statistical analysis

We applied the PCC to quantify the association between each individuals’ scores on specific topic and each individual’s rs10455872 status, for each learned topic. PCC measures the strength of a linear association between two variables. PCC also can generate a correlation coefficient denoted by *r ∈* [– 1,1], which shows the direction of the correlation.

We used the individual-topic matrix, *W ∈ R*^*n × k*^, generated by NMF to calculate the PCC with the genetic variants. Each column vector of *W*(:,*j*) of the matrix *W* represented a topic vector with scores on all the individuals. We used each column vector as the predictor variable *x* and the number of minor alleles (0, 1, or 2) at rs10455872 of each patient as variable *y.*

## Supporting Information

Table S1. Results with *topic k=10, 20,30*

